# Scaling of anesthesia-dependent cerebrospinal fluid dynamics across rat and pig brains

**DOI:** 10.64898/2026.07.10.737206

**Authors:** Natalie Beschorner, Miriam L. Navarro, Marko Rosenholm, Björn Sigurdsson, Nakul Raval, Emily Beaman, Sara Marie Ulv Larsen, Louise Møller Jørgensen, Clara Madsen, Visti H. Stenmo, Gerda Thomsen, Kristoffer Brendstrup-Brix, Claus Svarer, Maiken Nedergaard, Gitte Moos Knudsen

**Affiliations:** Center for Translational Neuromedicine, University of Copenhagen, Copenhagen, Denmark; Department of Neuroscience, Faculty of Health and Medical Sciences, University of Copenhagen, Copenhagen, Denmark; Neurobiology Research Unit, Copenhagen University Hospital Rigshospitalet, Copenhagen, Denmark; Division of Pharmacology and Pharmacotherapy, Faculty of Pharmacy, University of Helsinki, Helsinki, Finland; Yale School of Medicine, Department of Psychiatry, New Haven, CT, USA; Department of Psychiatry and Behavioral Sciences, University of Washington School of Medicine; Seattle, WA, USA; Copenhagen Spine Ressearch Unit, Copenhagen University Hospital Rigshospitalet-Glostrup, Copenhagen, Denmark; Department of Clinical Medicine, University of Copenhagen, Copenhagen, Denmark; Center for Translational Neuromedicine, University of Rochester Medical Center, Rochester, NY, 14642 USA

**Keywords:** CSF dynamics, pig, porcine model, gyrencephalic brain, cisterna magna cannulation, neuroimaging, glymphatic system, propofol, ketamine/dexmedetomidine, [^99m^Tc]-DTPA, SPECT

## Abstract

This study presents a novel *in vivo* neuroimaging approach using dynamic single photon emission computed tomography (SPECT/CT) to investigate cerebrospinal fluid (CSF) dynamics in pigs, a translationally relevant model due to their human-like brain structure. The distribution, brain penetration of [99mTc]-DTPA and subsequent clearance were followed by brain SPECT for three hours after injection into the cisterna magna of anesthetized pigs and rats. To investigate the effects of anesthesia and across-species effects, we examine CSF dynamics under two types of anesthesia, propofol and ketamine/dexmedetomidine (K/D), and compare the outcome in pigs to that of rats, in which we also compared isoflurane.

Propofol and K/D produced largely similar tracer distribution patterns across both pigs and rats: In both species, K/D was associated with higher tracer penetration into the dorsal striatum compared to propofol while neither species showed a tracer accumulation difference in the thalamus. K/D also increased intracranial radiotracer retention and reduced urinary tracer clearance in rats, but not in pigs. In rats, propofol and isoflurane showed similar tracer distribution, reflecting their shared GABAergic mechanism of action. The differences observed between pigs and rats may reflect species-specific physiology, differences in anesthesia dosing, or methodological factors.

The work demonstrates the feasibility of using SPECT/CT to study CSF transport in the large gyrencephalic pig brain to advance understanding of human brain fluid dynamics.

## Introduction

Traditionally, cerebrospinal fluid (CSF) has been viewed primarily as a passive component of the central nervous system, providing mechanical protection and support to the brain. Over the past decade, however, growing evidence has highlighted a more active role of CSF in maintaining brain homeostasis through the distribution of energy metabolites^1^, signaling molecules^2^ and the clearance of protein waste^3^. Early studies from the 1970s proposed that CSF enters the brain along perivascular spaces, exchanges with interstitial fluid (ISF), and thereby contributes to fluid and solute transport within the brain parenchyma^1–3^. Renewed interest in this concept led to the description of the “glymphatic system,” a brain-wide perivascular transport pathway characterized by periarterial CSF influx, exchange with ISF, and perivenous efflux, with astrocytic AQP4 water channels playing a central role in facilitating fluid movement^4,5^. The term glymphatic was introduced based on the involvement of glial cells and the observation that this pathway supports the clearance of amyloid-β and other metabolites from the brain imitating the lymphatic system. Downstream drainage is thought to occur via meningeal and peripheral lymphatic routes, linking central nervous system (CNS) waste removal to systemic circulation^6–8^. The activity of the glymphatic system has been shown to depend on brain state, making the choice of anaesthetic a critical experimental consideration. In rodents, tracer influx from the CSF into the brain parenchyma is greatest during natural sleep and in sleep-like states induced by ketamine/xylazine or dexmedetomidine (DEX) anesthesia, whereas it is markedly reduced during wakefulness and under isoflurane anesthesia^9–17^. Together, these studies have established CSF circulation and perivascular transport as important contributors to brain homeostasis. Dysfunction of these pathways has been associated with several neurological disorders, including Alzheimer’s disease, hydrocephalus, traumatic brain injury, cerebral small vessel disease, and stroke^4,18,19^.

Despite major advances, many aspects of glymphatic function remain under active investigation. Most experimental evidence originates from rodent studies, which enable invasive and high-resolution approaches. The rodent brain does, however, differ substantially from the human brain in anatomy, physiology, and sleep-wake organization^20,21^. Although a multitude of clinical studies have supported the existence of CSF clearing pathways in the human brain^22–25^, the limited feasibility of invasive measurements in humans has constrained mechanistic investigations. Large-animal models offer an important opportunity to bridge this translational gap. In particular, the pig brain shares key anatomical and physiological features with the human brain, including gyrencephaly, a high white-to-gray matter ratio, large brain size, and diurnal sleep-wake patterns^21,26–29^, making the pig a highly relevant translational model for studying brain fluid dynamics and clearance mechanisms. Accordingly, the domestic pig has become an increasingly important model in neuroscience, with well-established applications in behaviour research^30,31^, pharmacology^32^, neurological disease models^30,33^ and advanced neuroimaging^34–37^. Until now, methods for dynamic quantitative *in vivo* assessment of CSF transport and clearance in large-animal models has been limited. This methodological gap has hindered mechanistic and longitudinal studies of brain fluid dynamics in translational models.

To address this cross-species gap, we here establish a translational pig model for quantitative investigation of CSF transport by combining Cisterna Magna (CM) cannulation with dynamic whole-brain single-photon emission computed tomography/computed tomography (SPECT/CT) with the radiotracer [99mTc]-DTPA. CM cannulation is one of the most widely used approaches to study glymphatic transport in rodents, providing a rapid, reliable means to introduce tracers into the CSF^15,16,38,39^. Bèchet et al. has previously adapted this technique from rodents to pigs using a surgical procedure, demonstrating extensive perivascular solute transport in the gyrencephalic brain via *ex vivo* fluorescent microscopy^29,40^. Here, we further adapt the CM cannulation in pigs by performing a direct percutaneous puncture using an epidural needle, avoiding open surgery with potential impact on glymphatic function but also reducing procedure duration, technical complexity, and invasiveness. We then use the platform to examine how anaesthetic regimens with either propofol and ketamine/dexmedetomidine (K/D) influence CSF tracer dynamics and compare the findings with corresponding observations in rats. Propofol is a common anesthetic in both pig and human surgery while in rodents, K/D has been shown to promote glymphatic activity^10,11,13–17^. We then compare findings with rats imaged with SPECT/CT modality where we also investigate effects of isoflurane, the established benchmark GABAergic anesthetic, to another GABA modulator, propofol, which has not previously been used in rodents in this context, and to K/D. This cross-species approach enables us to assess whether anesthesia-dependent CSF transport and clearance are conserved across mammalian brains that differ substantially in size and anatomical complexity.

## Methods

### Experiments performed in pigs

#### Animals, housing and anesthesia

All animal experiments were performed in accordance with the European Communities Council Resolves of 22 September 2010 (2010/63/EU), approved by the Danish Veterinary and Food Administration’s Council for Animal Experimentation (Licence No. 2022-15-0201-01156), and reported in accordance with the ARRIVE guidelines.

Ten female pigs (Landrace x Yorkshire x Duroc crossbreed, aged 10 weeks and weighing about 20 kg), were obtained from a local breeder. Upon arrival at the Department of Experimental Medicine, University of Copenhagen, the animals were examined by a veterinarian and acclimatized to their new surrounding for one week. The pigs were housed in pens (3.8–5.4 m^2^) under controlled environmental conditions (relative humidity 45–70 %, ambient temperature 19–23 °C, and a 12h light-/-12-h dark cycle) with straw bedding and environmental enrichment. Animals had free access to drinking water, were fed twice daily, and were fasted 16 h before anesthesia on the experimental day. All experiments were conducted during the light phase, the active of the animals, equivalent to the awake phase for pigs.

On the experimental day, pigs were premedicated with an intramuscular injection of 0.14 mL/kg Zoletil® 50 Vet (Virbac, Kolding, Denmark) mixture, weighed and intubated. Each animal was fitted with a bladder catheter, a peripheral venous catheter on the right ear, a percutaneous femoral artery catheter (Arrow International Inc, Reading, PA, USA) and a catheter inserted through a small incision into the right mammary vein. All incisions sites were infilatrated subcutaneously with a mixture of 10 mg/mL xylocaine (AstraZeneca A/S, Copenhagen, Denmark) and 5 mg/mL Bupivacaine (Amgros I/S, Copenhagen, Denmark).

Two anesthesia regimes were evaluated: 1) PRO group (n=5) receiving maintainance anesthesia with an intravenous infusion of 12.5 mg/kg/h propofol (Fresenius Kabi AS, Halden, Norway) and 5 µg/kg/h fentanyl; and 2) K/D group (n=5) receiving 14 mg/kg/h ketamine together with dexmedetomidine (DEX), administered initially as an intramuscular injection of variable doses (0.018-0.02 mg/kg) before catheter placement followed by maintainance anesthesia with intravenous dexmedetomidine (0.008-0.016 mg/kg/h).

After a short transfer by trolley to the SPECT imaging facility, the animals were connected to a ventilator with 34 % O_2_, and received a slow intravenous infusion of isotonic saline and glucose, adjusted according to the physiological needs. During SPECT acquisition, the pigs were continuously monitored by visual assessment, including blink reflexes, and by manual and digital recording of physiological parameters. These included arterial blood pressure (through the femoral arterial catheter), heart rate (HR) and peripheral oxygen saturation (SpO_2_; measured with a pulse oximeter placed on the tail), respiratory rate and tidal volume (controlled by the ventilator), end-tidal carbon dioxide (EtCO_2_; measured using a capnograph positioned between the endotracheal tube and the ventilator circuit), and rectal temperature. Most of the physiological data were digitally recorded using VitalRecorder^41^ software running on a laptop connected to a Intelli Vue Phillips monitor. For each pig, physiological variables were summarized as mean values across the first 3h scanning period included in this article (from the total 6h of scanning) (Table 1).

**Table 1.** Overview of pigs.

| Pig no. | Group | Weight | Body position | MAP | Heart Rate | Resp. volume | Resp. /min |
| --- | --- | --- | --- | --- | --- | --- | --- |
| 1 | PRO | 20 | Left lateral | 57.4 ± 6.5 | 75.3 ± 5.1 | 200 ± 0 | 14.82 ± 0.4 |
| 2 | PRO | 21 | Left lateral | 82.2 ± 9.2 | 95.1 ± 11.8 | 250 ± 0 | 20 ± 0 |
| 3 | PRO | 19 | Left lateral | 65.9 ± 10.8 | 77.0 ± 3.5 | 200 ± 0 | 20.6 ± 0.9 |
| 4 | PRO | 24 | Left lateral | 75.3 ± 5.4 | 107.3 ± 4.5 | 270 ± 0 | 22 ± 0 |
| 5 | PRO | 19 | Left lateral | 74.0 ± 20.3 | 84.8 ± 11.2 | 201.6 ± 11.7 | 18.7 ± 0.8 |
| 1 | K/D | 20 | Left lateral | 66.5 ± 0.76 | 71.6 ± 5.5 | 131.2 ± 15.5 | 17.5 ± 0.5 |
| 2 | K/D | 22 | Left lateral | 90.6 ± 12.2 | 89.5 ± 8.8 | 220 ± 0 | 17 ± 0 |
| 3 | K/D | 22 | Left lateral | 113.1 ± 7.3 | 71.0 ± 5.3 | 230 ± 0 | 19 ± 0 |
| 4 | K/D | 9 | Left lateral | 107.9 ± 3.5 | 49.6 ± 2.9 | 210 ± 0 | 16 ± 0 |
| 5 | K/D | 22 | Left lateral | 92.6 ± 6.9 | 84.2 ± 34.3 | 190 ± 23.1 | 19.2 ± 1.0 |

Immediately after completion of the final scan, the pigs were euthanized by intraveneous administration of an overdose of 20 mL sodium pentobarbital (ScanVet Animal Health A/S, Fredensborg, Denmark).

#### Cisterna magna cannulation

The direct percutaneous approach to the cisterna magna (CM) is illustrated in Figure 1b. In one animal, CM cannulation was unsuccessful, resulting in inadvertent injection of the radiotracer into the neck musculature (Supplementary Figure 1). This animal was excluded from further analysis.

**Figure 1.**
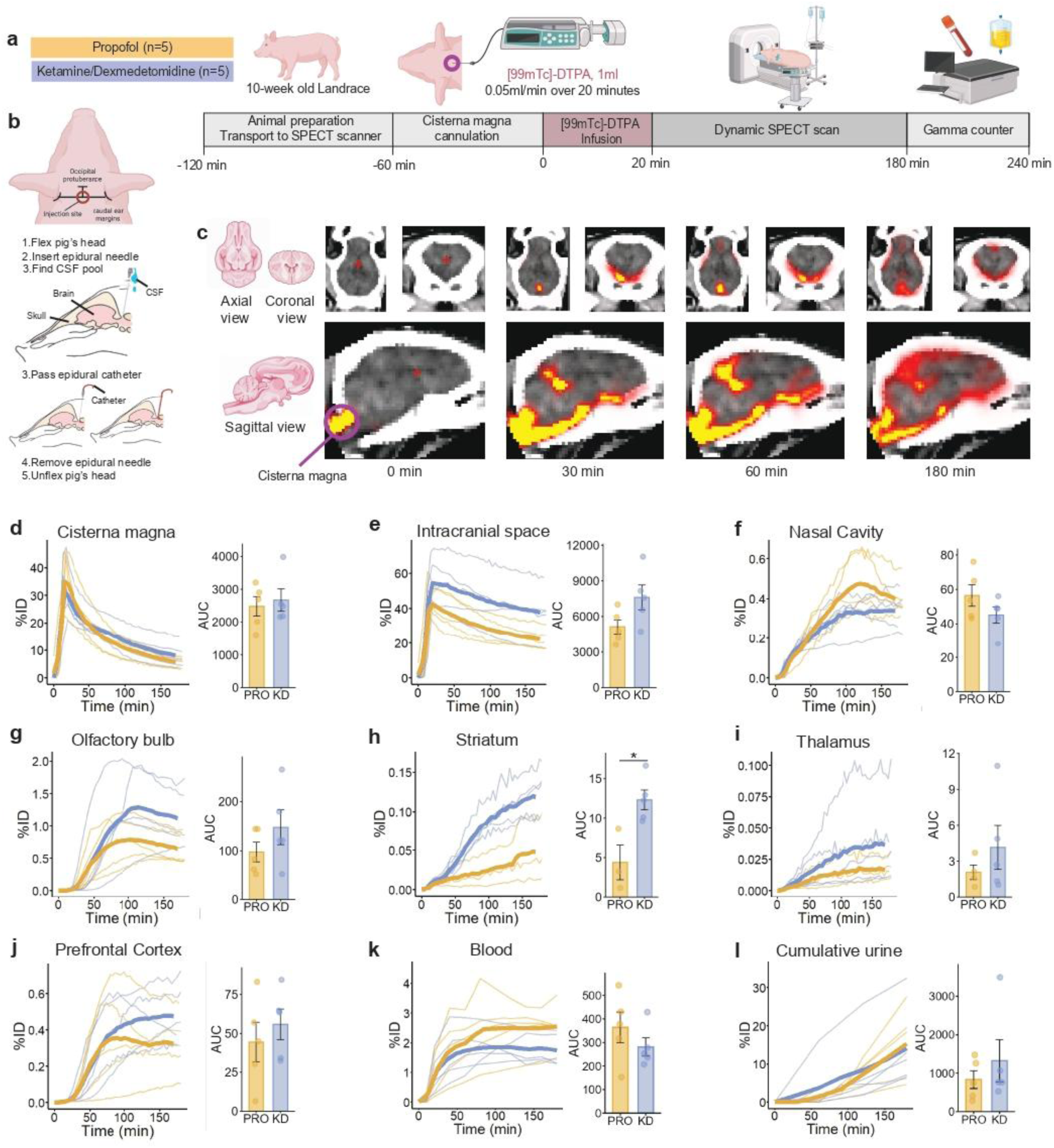
Experimental design and dynamic SPECT/CT assessment of CSF tracer distribution in pigs under different anesthesia regimens. (a) Study design comparing propofol (PRO) and ketamine/dexmedetomidine (K/D) anesthesia in 10-week-old Landrace pigs (n=5 per group), including cisterna magna (CM) cannulation, tracer infusion, and dynamic SPECT/CT acquisition._Experimental timeline and procedural workflow from animal preparation to imaging and post-scan gamma counting. (b) CM cannulation steps. (c) Representative SPECT/CT images showing tracer distribution over time (0, 30, 60, and 180 min) in axial, coronal, and sagittal views, illustrating tracer entry from the CM and subsequent brain-wide dispersion. A red cross in the middle of the brain marks the location of the SPECT/CT slices shown. (d-j) Time-activity curves (TACs, % injected dose) and corresponding area under the curve (AUC) analyses for CSF and brain regions, including CM (d), intracranial space (e), nasal cavity (f), olfactory bulb (g), dorsal striatum (PRO group n=3) (h), and thalamus (PRO group n=4) (i), and prefrontal cortex (j) comparing PRO (red) and K/D (blue) groups. Significant stars are, * if p-value < 0.05, ** p-value is <0.01,***p-value <0.001. (k-l) Additional systemic tracer distribution and clearance dynamics of radioactivity TACs and AUC for blood (k), and cumulative urinary tracer excretion over time (l).

CM cannulation was performed using a Perifix® Soft Tip 701 filter sets (B Braun, Melsungen, Germany). The pig was placed in the left lateral decubitus position with the neck flexed by an assistant. The insertion point was located at the midpoint of an imaginary line connecting the caudal margins of the ears, immediately caudal to the occipital protuberance (Figure 1b). The needle was advanced with the stylet in place through approximately 5-7 cm of neck musculature towards the CM. Occasionally, a subtle loss of resistance was appreciated as the meninges were penetrated. -The stylet was then removed, and free flow of CSF from the needle hub confirmed correct entry into the subarachnoid space. If bone was encoundted during needle advancement, the needle was withdrawn and redirected slightly before a new attempt was made.

Following confirmation of intrathecal needle placement, the epidural catheter was advanced though the needle several centimeters beyond the needle tip into the subarachnoid space. While maintaining the catheter in fixed position, the needle was carefully withdrawn over the catheter to minimize the risk of catheter displacement. The catheter was then controlled for free flow of CSF and temporarily secured to the skin with adhesive tap. Catheter position in CM was then verified by CT. If the catheter extended to far intracranial within the subarachnoid space, it was withdrawn slightly and CT imaging was repeated until satisfactory position in CM was confirmed. Once the final position had been verified, the catheter was permanently secured with skin sutures and adhesive tape.

#### SPECT scanning protocol

All pigs were placed in left lateral position. Because the full body could not be covered by the scanner’s field of view, we positioned the head and upper cervical spine within the field of view. Subsequent scans confirmed the capture of close to 100% of the injected activity. All pigs underwent a dynamic SPECT-scanning with the high-resolution AnyScan TRIO® SPECT/CT imaging system (Mediso Medical Imaging Systems, Budapest, Hungary). A detailed SPECT scanning protocol is included in the Supplementary Material. [^99m^Tc]-DTPA (12.5 mg/mL, TechneScan DTPA, Curium Pharma; MW 489 Da) was prepared at the Copenhagen University Hospital, Rigshospitalet. Right before the start of the emission scan, a high-resolution CT scan was performed. We use [^99m^Tc]-DTPA as a radiotracer due to its confinement to the CSF compartment and its inability to cross an intact blood–brain barrier, allowing clear assessment of CSF distribution and clearance pathways^38,42,43^. Afterwards, 1-2 mL of [^99m^Tc]-DTPA was slowly injected over 20 min at a rate of 0.05 - 0.1 mL/min to avoid a drastic increase of intracranial pressure. The individual injected dose was calculated for each pig by measuring the activity of the vial with a scintillation counter before and after tracer injection subtracting background activity and the activity left in the tubing after the scan. Data acquisition lasted a total of ∼3h, including 20 minutes of tracer injection. The resulting voxel size was 1.8×1.8×11.8 mm and each frame lasted approximately 6 min. The SPECT-scanner with the described set-up was quantitatively calibrated for Tc-99m. Values in the reconstructed SPECT-images were therefore saved in Bq/mL. More details are available in the Supplementary Material.

#### SPECT/CT processing and analysis

SPECT processing steps included: 1) conversion of the DICOM format using MedisoConvert in MATLAB version 9.14.0.2239454 (R2023a) (in-house code). 2) Manual co-registration of the MRI pig atlas^44^ to the CT and SPECT performed with an in-house program called using a 12-parameter transformation in MATLAB version 8.3.0.532 (R2014a). 3) The prefrontal cortex, olfactory bulbs, dorsal striatum and thalamus were obtained from a MRI atlas^44^ co-registered to the SPECT image. We also manually delineated regions of interest (ROIs) using ITK-SNAP (version 4.0.1)^45^. Manual ROIs included: intracranial space (including brain tissue and cranial CSF spaces), nasal cavity, eyes and cisterna magna. 4) Extraction of the time activity curves (TACs) from dynamic SPECT images for further analysis using an in-house code in MATLAB version 9.14.0.2239454 (R2023a). TACs in Bq/mL were multiply by the volume of the ROI and divided by the injected dose to obtain percentage of injected dose (%ID) used to visualize radioactivity over time. TACs for each ROI are presented in line plots in Figure 1d-l and 3d-l individually for each pig in thin lines plus the mean in a thick line and coloured by anesthesia group.

Area under the curve (AUC) of each TAC was calculated using the trapezoidal rule implemented in the *trapz* function from the pracma package in R (version 4.6.0)^46^. AUC data is presented in column graphs with individual points for each pig (Figure 1).

To reduce MRI atlas-defined ROIs being affected by partial volume spillover from adjacent CSF spaces, the deeper brain regions striatum and thalamus were selected for the primary ROI analyses. However, for dorsal striatal TACs and for one thalamic TAC, the signal displayed a profile resembling CSF-like activity (the TAC shape was similar to the intracranial space TAC, which was considered a surrogate of CSF activity) suggesting slight coregistration inaccuracies. A plot displaying all striatal and thalamic TACs normalized to their maximum activity is provided in Supplementary Figure 2. This figure illustrates the differences in TAC shapes for striatum and thalamus ROIs as well as their similarities to brain and intracranial space TACs (intracranial space activity will be driven mostly by the CSF activity, therefore CSF-like TACs). Pigs with CSF-like TAC shapes in teh thalamus and striatum were removed from TAC plots and AUC comparison from Figure 1h, i and Figure 2h, i.

**Figure 2.**
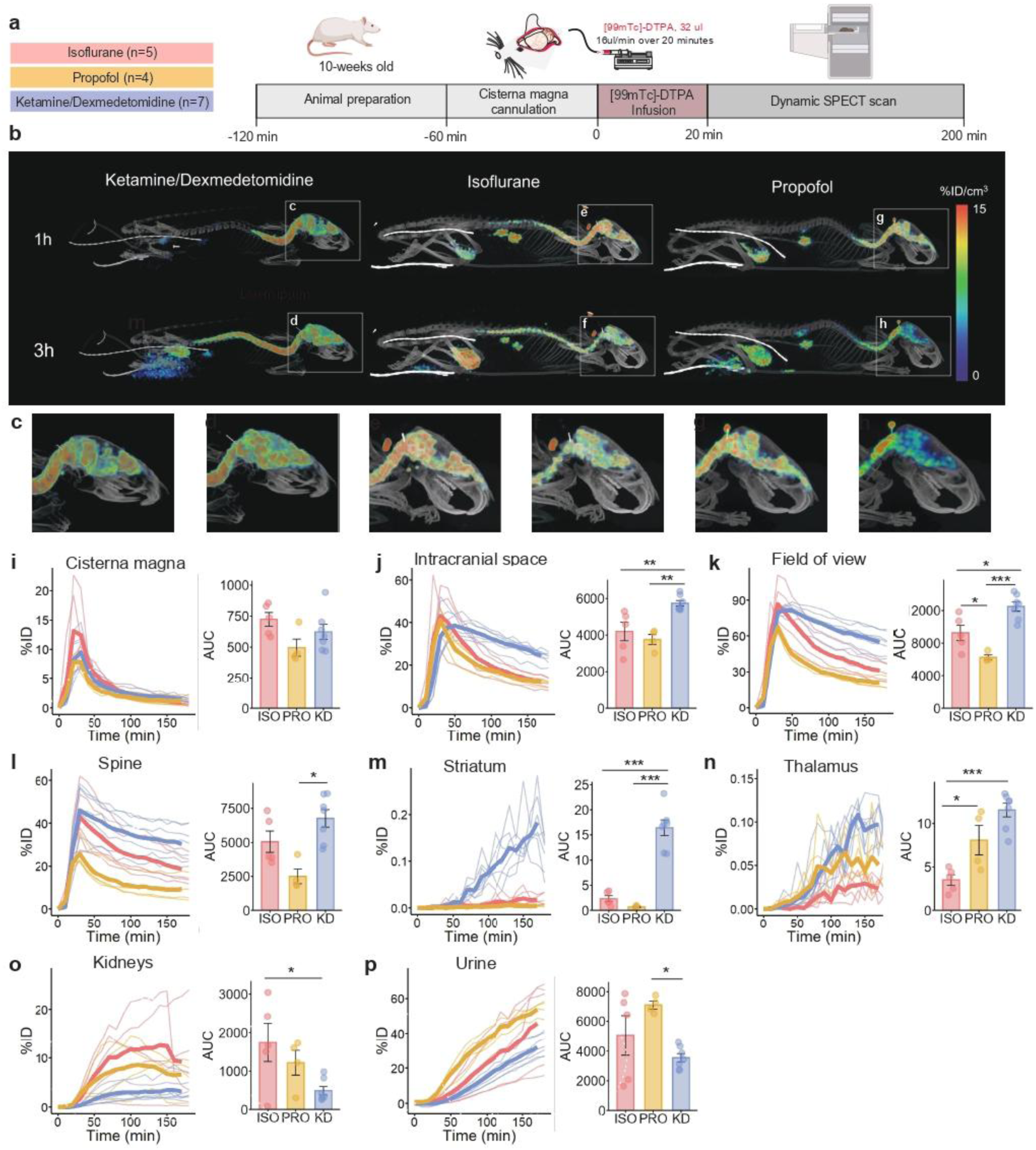
Anesthesia-Dependent CSF Tracer Dynamics in Rats. (a) Study design comparing propofol (PRO), ketamine/dexmedetomidine (K/D) and isoflurante (ISO) anesthesia in 10-weeks old rats, including cisterna magna (CM) cannulation, tracer infusion, and dynamic SPECT/CT acquisition. Experimental timeline and procedural workflow from animal preparation to imaging. (b) and (c) Representative SPECT/CT images showing tracer distribution over time. (i-p) Time-activity curves (TACs, % injected dose) shown individually for each rat in thin lines plus the mean in a thick line and coloured by anesthesia group, and their corresponding area under the curve (AUC) analyses for different ROIs, including CM (i), intracranial space (j), field of view (k), spine (l), striatum (m), and thalamus (n), comparing PRO (red) and K/D (blue) groups. No value is given if p-value >0.01; if p-value is <0.05 = “*”; if p-value is < 0.01 = “**”; and if p-value is <0.005 = “***”. (o-p) Additional systemic tracer distribution and clearance dynamics of radioactivity TACs and AUC for kidneys (o), and cumulative urinary tracer excretion over time (p).

#### Blood and urine sampling

Arterial blood samples of 2 mL each were manually drawn at 0, 5, 10, 20, 30, 40, 60, 80, 100, 120, 140, 160, and 180 min after start of the scan. Urine samples were collected every 30-60 min (if there was urine produced in the collection bag) from the urinary bladder catheter and the volume was noted after which the urine bag was emptied. Radioactivity in whole blood and urine samples was measured with an automated gamma-counter (Hidex AMG, Hidex Oy, Turku, Finland) at the end of the experiment and decay corrected to the start of the scanning. To calculate %ID for blood radioactivity, we assumed 65 mL of blood per kg and used the average pig’s weight of 20 kg, resulting in 1300 mL of blood in a pig. For each blood sampling timepoint, the estimated blood volume was multiplied with the blood radioactivity concentration (Bq/mL normalized to the injected dose) to obtain total %ID in blood as a function of time. To obtain cumulative urine radioactivity excretion over time, we multipied the urine radioactivity concentration (Bq/mL) by urine volume produced and summed over time. TACs and AUC are calculated plotted in Figure 1d-l and 3d-l.

#### Statistical analysis

Comparisons of AUC values between the two anesthesia groups (propofol and K/D) were performed separately for each ROI using independent two-sample t-tests without correcting for multiple comparisons due tot he exploratory nature of the study. For each comparison, mean group differences, 95% confidence intervals, and p-values were calculated. This analysis was implemented in R using the t.test, dplyr, and broom packages. All statistical tests were two-tailed. Statistical significance was set at 0.05. Statistical analyses were performed in R (version 4.3.1; R Core Team, 2024). No statistical comparisons of physiological parameters were performed between groups.

### Experiments performed in rats

#### Animals and housing

All rodent experiments were approved by the Danish Animal Experiments Inspectorate and carried out at the Center for Translational Neuromedicine, University of Copenhagen, in accordance with the European Directive 2010/63/EU under the license 2020-15-0201-00581 and followed the ARRIVE guidelines. In total, 17 female 10 weeks old Sprague-Dawley rats (ISO n=5, PRO n=5, K/D n=7, 200-250 g, Janvier, Le Genest-Saint-Isle, France) were used for [^99m^Tc]-DTPA experiments. One of the five rats (PRO group) died during the scan and was therefore excluded from the analysis. K/D rats data has been previously published in Lilius *et al*. 2022 as the control group data^47^. Rats were housed in pairs in individually ventilated plastic cages with ad-libitum access to water, environmental enrichment and a standard rodent diet in temperature-controlled rooms with a 12h light/dark cycle. All experiments reported were performed during the light phase of the animals, equivalent to the sleep stage for rats.

#### Cisterna Magna (CM) cannulation and anesthesia

The CM cannulation was performed as previously described^39^. For induction of anesthesia, rats were anesthetized with 3-4% of isoflurane. At the beginning of the surgery, lidocaine (1 mg/kg) was administered. During the surgery, anesthesia was maintained with 2-3% isoflurane. After loss of pain response, animals were placed in a stereotaxic frame with the neck flexed at 30–40°. The atlanto-occipital dura overlying the CM was exposed by carefully removing and pushing apart the neck muscles. A 30G needle connected to a PE10 tubing (Scandidact) was carefully inserted into the intrathecal space, kept in place with cyanoacrylate glue (Loctite) followed by application of dental cement. For placement of the intravenous catheter, the tail vein was dilated by gentle application of a heat pad (Thermopad). A 24G intravenous cannula (BD Neoflon) was inserted in the right lateral tail vein and fixed in place with surgical tape. Afterwards, the rats were transferred to the imaging bed.

Three anesthetic regimens were used: isoflurane, propofol, and ketamine/dexmedetomidine. For isoflurane anesthesia, anesthesia was induced with 3–4% isoflurane and maintained with 1.5–2% isoflurane in 100% oxygen. For propofol anesthesia, animals were initially anesthetized with isoflurane for CM cannulation, Propofol infusion was then initiated at 40 mg/kg/h via the venous catheter while isoflurane was gradually discontinued over 5 min. For ketamine/dexmedetomidine anesthesia, ketamine (100 mg/kg) and dexmedetomidine (0.5 mg/kg) were administered together by subcutaneous injection before tracer injection.

Body temperature and respiratory rate were continuously monitored throughout the experiment (Table 2). Directly after SPECT/CT scanning, the rats were killed by cervical dislocation.

**Table 2.** Overview of rats.

| Rat no. | Group | Weight (g) | Injected Dose (MBq) | Rat position | Respiration /min |
| --- | --- | --- | --- | --- | --- |
| 1 | PRO | 230 | 15.6 | prone | 64.2 ± 2.1 |
| 2 | PRO | 220 | 8.9 | prone | 61.3 ± 4.0 |
| 3 | PRO | 230 | 13.6 | prone | 55.1 ± 7.9 |
| 4 | PRO | 230 | 11.7 | prone | 59.5 ± 3.2 |
| 1 | ISO | 230 | 9.2 | prone | 31.8 ± 3.9 |
| 2 | ISO | 220 | 6.1 | prone | 37.1 ± 3.5 |
| 3 | ISO | 240 | 20.1 | prone | 42.7 ± 2.8 |
| 4 | ISO | 230 | 7.9 | prone | 55.1 ± 7.2 |
| 5 | ISO | 230 | 14.5 | prone | 58.4 ± 9.4 |
| 1 | K/D | 235 | 10.4 | prone | 78.5 ± 8.3 |
| 2 | K/D | 230 | 20.7 | prone | 83.5 ± 10.2 |
| 3 | K/D | 230 | 22.3 | prone | 65.6 ± 7.9 |
| 4 | K/D | 220 | 25.6 | prone | 63.9 ± 8.6 |
| 5 | K/D | 220 | 13.5 | prone | 67.5 ± 7.2 |
| 6 | K/D | 230 | 16.7 | prone | 71.3 ± 6.4 |
| 7 | K/D | 230 | 19.3 | prone | 81.0 ± 9.6 |

#### SPECT/CT scanning protocol

Dynamic whole-body SPECT/CT scanning on rats in prone position was performed with the Vector4CT (MILabs, Utrecht, Netherlands) system. Images were acquired with a high energy ultra-high-resolution rat 1.8 mm pinhole collimator (HE-UHR-RM 1.8 mm diameter) with a spatial resolution of approximately 0.9 mm. Total data acquisition time was 220 minutes with twenty-two frames at 10 minutes per frame (only the first 180 min are included in the analysis). The temperature of each animal was monitored and maintained with a thermostatically regulated heating pad. Subsequently to the SPECT scan, a whole-body CT image was acquired, co-registered to the SPECT image and used as an anatomical reference. The full-body CT scan was acquired with 55 kV tube voltage, 75 ms exposure time, x11 binning and 360 projections with a steep angle of 0.75 degrees over 9 minutes. [^99m^Tc]-DTPA (12.5 mg/mL, TechneScan DTPA, Curium Pharma; MW 489 Da) was used as a radioactive tracer for imaging. For intracisternal administration, 32ul of [^99m^Tc]-DTPA at a speed of 1.6 μL/min was infused with a Hamilton Gastight 1700 syringe in a micro infusion pump. Radioactivity ranged between 10-30 MBq and was measured with a VIK-202 dose calibrator (Comecer).

#### SPECT analysis

Images acquired were reconstructed with Similarity-Regulated Ordered Subsets Estimation Maximization (SROSEM) using a voxel size of 300 μm and five iterations with correction for attenuation and tracer decay in MILabs Reconstruction software 10.16 (MILabs, Utrecht, Netherlands). ROIs were manually drawn in ITK-SNAP software^31^ using the acquired SPECT and CT images. ROIs in the central nervous system included the intracranial space, the CM, the striatum and the thalamus. ROIs that are relevant to CSF clearance routes included the head and the urinary system (left and right kindey, bladder and urine collected on the pad placed under the animal). The total injected activity in each ROI was calculated as the percentage of the injected dose (%ID) using MATLAB R2019a (Mathworks).

#### Statistical analysis

The AUC from each TAC (for 3 scanning hours) was calculated using the trapezoidal rule implemented in the *trapz* function from the pracma package in R^46^. To compare between propofol, isoflurane and K/D groups, an ANOVA was fitted with group and post hoc pairwise group comparisons and p-values were adjusted with Tukey. This analysis was implemented in R using the aov, emmeans, and broom packages. All statistical tests were two-tailed. Statistical significance was set at 0.05. Statistical analyses were performed in R (version 4.6.0; R Core Team, 2024).

## Results

### Tracer distribution in pigs

A diagram of the experiments and groups is shown in Figure 1a. First, we evaluated whether SPECT imaging of CSF flow in the pig brain revealed dynamic tracer distribution patterns comparable to those described in rodents ^32^. During the 180-minute dynamic scan, the injected tracer gradually dispersed from the CM, first along ventral pathways near the circle of Willis, around the cerebellum and then dorsally. Tracer penetration into deeper brain structures was limited (Figure 1c).

[^99m^Tc]-DTPA tracer activity in the CM reached its maximum shortly after infusion stopped and gradually decreased over time as tracer left the CM and distributed to the intracranial space and the spinal cord (Figure 1d). Intracranial tracer activity peaked immediately after infusion, with variable peak levels across animals ranging from 25% to 65% of the total injected dose. While there were differences in peak activity, the TACs declined at a similar rate in all experiments. By the end of the 180-min scans, intracranial space values had decreased by about 10-20% of their peak values (Figure 1e). In the nasal cavity ROI (a possible CSF efflux pathway), most TACs showed a gradual increase until plateaued between 1-2 hours post-injection (Figure 1f). The olfactory bulb showed a TAC shape similar to the nasal cavity (Figure 1g), with the distinction that tracer concentration tended to be higher under K/D in the olfactory bulb, while propofol was associated with slightly higher concentrations in the nasal cavity, but no significant differences were observed in AUCs between propofol and K/D anesthesias in these ROIs (CM, intracranial space, nasal cavity, and olfactory bulb difference in AUC estimates between K/D and propofol with [95% CI] = 190 [-853, 1233], 2503 [-407, 5413], - 11.4 [-29.6, 6.8], and 50 [-48, 148]; p-values = 0.7, 0.08, 0.2, and 0.3, respectively).

[^99m^Tc]-DTPA tracer penetrance into the brain was assessed in the prefrontal cortex, thalamus and dorsal striatum. The prefrontal cortex showed a fast increase at 1h post injection after which it plateaued or increased slowly (Figure 1j). In the thalamus and striatum the radioactivity steadily increased over time, reaching a maximum of 0.05-0.15 %ID by the end of 3 hours (Figure 1h, i). The thalamus and striatum represent deeper brain regions, indicating tracer influx into subcortical areas. The AUC showed higher tracer brain influx under K/D compared to propofol for the striatum (estimate difference = 8.9 [4.7, 13.2]; p-value = 0.04), but no difference in the prefrontal cortex nor the thalamus (estimate difference = 11.4 [-26.1, 48.8] and 0.77 [-4.6, 6.2]; p-value = 0.5 and 0.3, respectively).

To assess potential asymmetries arising from the pig being positioned on its left side, we compared radiotracer AUC between the right and left eye, prefrontal cortex, and striatum. Overall, there was a general higher proportion of tracer in the right hemisphere for the propofol group on the three ROIs (mean ratio right to left 1.4), whereas K/D group showed higher radioactivity in the right hemisphere for the prefrontal cortex (mean ratio 1.2) but lower for the eyes and striatum ROIs (mean ratio for both ROIs 0.7). The asymmetries could come from body positioning or catheter placement.

### CNS tracer efflux in pigs

To evaluate tracer efflux from the CNS, we measured radioactivity in the blood and urine during the scan. Blood radioactivity was first detected 10 minutes after tracer injection, and steadily increased to reach levels of 2-3 %ID (Figure 1k). Blood radioactivity peaked for the K/D group at around 1 hour; in the propofol group, the peak occurred about 2.5 hours after injection but there was no difference in blood AUC between groups. Urine radioactivity increased steadily across the entire 3h scanning to reach its maximum around 10% of ID (Figure 1l). Despite that the urine cumulative radioactivity AUC did not differ between the PRO group (9.1± 5.8 %ID) and the K/D group (14.1 ± 9.5 %ID), urine production at the end of the 3h was larger in K/D group (mean ± SD: 820 ± 127 mL) compared to PRO (340 ± 122 mL), suggesting lower radioactivity concentration per mL of urine in the K/D group. Blood and urine measurements revealed no differences in CNS tracer excretion between anesthesia conditions (blood AUC estimate difference = 14.8 [-181, 211], p-value = 0.8, and urine AUC estimate difference = - 1599 [-6781, 3581], p-value = 0.5).

### Tracer distribution and physiology in rats

Table 2 displays vitals in the rats and a diagram of the experiments and groups is shown in Figure 2a. TACs are presented in line plots in Figure 2i-p. At 60 minutes and 200 minutes post-injection, the highest tracer concentration was observed in the ventral brain (also in the olfactory bulb and the nasal cavity), CM region, and upper cervical spine (Figure 2b). A time-series of tracer distribution from 60 and 180 minutes is shown in Figure 2b-h.

Tracer activity in the intracranial space peaked at 40 %ID within 20 minutes of infusion started and was halved at the end of the 3-hr scan. Activity in the CM and in the head region followed similar trends, peaking at 65 %ID and 8 %ID, respectively, before declining. Possibly due to the shorter diffusion distances in the rats, the striatum of the rats showed significantly higher tracer retention (2 %ID at 220 minutes) compared to pigs (0.015 %ID at 240 minutes).

### Effects of anesthesia on tracer distribution in rats

Tracer dynamics collected under propofol anaesthesia did not differ from those collected under isoflurane anesthesia. For both types of anesthesia, however, the intracranial space and spine tracer retention was lower than for K/D anaesthesia (Figure 2j,k,l) (AUC mean difference between PRO and K/D with [95% CI] for intracranial space, field of view and spine ROIs respectively are -1982 [-3199, -765], -6252 [-8870, -3635], and -4270 [-6918, -1622]; and adj. p-values = 0.002, 7e-5, 0.002). K/D was also associated with higher tracer retention in the striatum compared to PRO and ISO (Figure 2m) (AUC mean difference between PRO and K/D = -15.7 [-20.5, -10.9], adj. p-value = 2e-6). Tracer influx in the thalamus was lower under isoflurane than under either K/D and PRO, whereas the latter two conditions did not differ significantly from each other (Figure 2n) (AUC mean difference between ISO and K/D, between ISO and PRO, and between PRO and K/D, respectively = -8 [-11.5, -4.5], -4.6 [-8.6, -0.5], and −3.4 [−7.2, 0.3]; adj. p-values = 0.0011, 0.02, and 0.07). We also detected faster clearance of the tracer from the CNS space (Figure 2k) (mean difference between PRO and K/D = -6252 [-8870, -3635]; and adj. p-values = 7e-5), into the periphery shown by increased urine radioactivity (Figure 2p) (mean difference between propofol and K/D= 3546 [661, 6431]; adj. p-values = 0.016).

## Discussion

In this study, we investigate how cerebrospinal fluid transport and clearance scale across mammalian brains that differ markedly in size and anatomical complexity. Using dynamic whole-brain SPECT/CT imaging following CM injection of [99mTc]-DTPA, we compare tracer distribution and clearance in pigs and rats, two species that differ by more than an order of magnitude in brain volume. We further examine whether anesthesia-dependent alterations in brain fluid dynamics are conserved across species by comparing the effects of propofol and ketamine/dexmedetomidine (K/D). While K/D increased tracer accumulation in the striatum in both species, its effects on intracranial tracer retention and systemic clearance differed slightly between pigs and rats. In pigs, intracranial retention showed a trend toward being higher and lower urine clearance under K/D like in rats, though these did not reach statistical significance. These findings suggest that some aspects of CSF transport are conserved across brain scales, whereas others may depend on species-specific physiology, experimental conditions, or scaling relationships that emerge in larger brains.

### Translational in vivo imaging of CSF dynamics in a large-animal model

Using post-mortem *ex vivo* imaging, Bèchet documented, for the first time, perivascular tracer influx in the porcine brain^29^. Unlike ex vivo approaches, SPECT imaging enables dynamic assessment of all brain regions, albeit with a lower spatial resolution, including deeper brain regions while the *ex vivo* approach can demonstrate cortical micro-distribution. Compared with MRI and contrast-based approaches, SPECT/CT enables longitudinal imaging with substantially lower tracer volumes, which is critical for preserving physiological intracranial pressure. Intrathecal contrast MRI studies are used to study CSF circulation in humans, but they are constrained by invasiveness and low time-scale sampling where the brain peak of radioactivity is usually missed^22,24^. The porcine gyrencephalic brain provides an important translational model that enables dynamic molecular neuroimaging with invasive CSF studies in an animal with a more human-relevant brain anatomy.

Building on Bèchet et al.^40^, we adapted CM cannulation for *in vivo* use by introducing a percutaneous approach that avoids neck muscle removal, thereby reducing surgical trauma and blood loss. Although we assured the correct catheter position with CT scan, subtle variations in the position of the tip of the catheter can introduce some variability in tracer distribution between the spinal and intracranial compartments or between left and right hemispheres. This may translate into interindividual differences in intracranial peak concentrations and left-right tracer asymmetries. Additional procedural factors, including subtle differences in effective tracer delivery (e.g. completeness of catheter flushing, reflux, preferential cranial versus caudal tracer distribution, leakage at the puncture site, or resistance during infusion), may also have contributed to the interindividual variability in peak intracranial tracer activity.To account for spatial and temporal differences in tracer delivery as measured in CSF, kinetic modeling could be useful. This would, however, require a more accurate brain segmentation not supported by the spatial resolution of the applied imaging techniques.

### Tracer distribution and clearance pathways

The radiotracer rapidly dispersed along the ventral brain surface and cerebellar sulci and subsequently penetrated into deeper brain regions and cleared via the nasal cavity, blood and urine. The radiotracer accumulated in the striatum and thalamus supporting that CSF-derived solutes can reach deep subcortical and cortical regions within hours, while the presence of nasal radioactivity supports the presence of olfactory CSF efflux pathways^48^. Due to the limited spatial resolution of the clinical SPECT scanner relative to the size of the pig brain, partial volume effects were clearly visible, particularly in the prefrontal cortex and olfactory bulbs.

### Effects of anesthesia across pigs and rats

Consistent with known DEX vascular effects^49^ and compared to propofol, K/D anesthesia was associated with higher mean arterial pressure and lower heart rate in the pigs. In both pigs and rats, K/D consistently increased radiotracer accumulation in dorsal striatum but not in thalamus. Methodological factors, including partial volume effects and ROI definition, are important considerations; however, TACs with CSF-like characteristics were identified and excluded prior to analysis. The remaining regional difference may therefore also reflect differences in ROI composition or regional anatomy. The dorsal striatum represents a relatively homogeneous anatomical structure with a well-defined vascular territory, whereas the thalamic ROI comprises multiple anatomically and physiologically distinct nuclei with different vascular territories. Averaging across this heterogeneous ROI may reduce sensitivity for detecting localized regional differences. We also refer to "striatal accumulation" rather than distinguishing influx from efflux, as the AUC analysis does not permit this distinction; kinetic modelling would be required to characterize directional tracer movement. Whether the observed regional differences reflect vascular architecture, perivascular transport, or other physiological factors remains to be determined.

In contrast to rats, we did not see statistical higher intracranial radiotracer retention in the pigs under K/D anesthesia. We speculate that this difference could be due a dose-dependent effect of DEX or variations in catheter positioning. In support of the former, as a post hoc observation, the two pigs receiving the higher DEX dose showed markedly greater CSF radiotracer retention than those receiving the lower dose, worth investigating in future work. We further speculate that the elevated intracranial CSF tracer concentration and reduced peripheral excretion observed under higher DEX doses in rats may reflect DEX-mediated reduction of CSF production, consistent with the known effects of other adrenergic blockers^50^. Finally, isoflurane and propofol anesthesia in the rats was associated with similar tracer distribution patterns supporting that both GABAergic anesthetics exert related effects on CSF transport.

### Limitations

Given the exploratory nature of this study, some findings should be interpreted cautiously; nonetheless, the results support the potential of *in vivo* glymphatic imaging in gyrencephalic brains. Future studies should involve assessment of automated ROI delineation and kinetic modeling of the tracer movements. A methodological limitation is the variability in catheter position, which can also affect the initial tracer distribution between the cranial and spinal CSF, shown as differences in the initial intracranial peak TAC.

### Conclusions

This study demonstrates the feasibility of dynamic SPECT/CT for investigating radiotracer based CSF transport and clearance in a large-animal model, the pig. In both pigs and rats, we found increased striatal K/D-related tracer influx, as also previously observed in mice. However, anesthesia-dependent differences in intracranial retention and CNS efflux were less consistent across species, in pigs, retention showed a trend toward being higher under K/D, though this did not reach statistical significance, this may reflect species-specific physiology or methodological factors such as dosing, ventilation, infusion protocols, body positioning, and differing circadian phase at the time of imaging (active phase in pigs vs. resting phase in rats). Overall, these findings highlight both the translational promise and current methodological limitations of large-animal glymphatic SPECT imaging, and emphasize the importance of standardized analysis strategies and future kinetic modeling across species.

