## Supplementary Materials and Figures for "Scaling of anesthesia-dependent cerebrospinal fluid dynamics across rat and pig brains"

**Supplementary Material**

*SPECT/CT-scan acquisition and reconstruction*

Scans were acquired on a Mediso Anyscan Trio SPECT/CT (Mediso, Budapest, Hungary) equipped with a 16-slice diagnostic CT and three gamma camera heads with 9.5 mm thick sodium iodide crystals mounted with multi-pinhole (MPH) brain collimators. Most relevant CT parameters were 120 kVp, 200 mAs, 1 s rota-tion time, 20 mm collimation, 1.25 mm slice thickness, 1.00 pitch and CT-brain table mode. Matrix size was 512x512 with a Field of View (FoV) of 500x500 mm. The axial FoV corresponded to the 17 cm axial FoV of the SPECT-scan, which was acquired with a 20% full width energy window at 140.5 keV, 1.14 zoom and 15 cm detector radius. The dynamic SPECT-scan consist-ed of a series of 36 frames or more. Each frame was acquired in 256x256 matrices with 2.13 mm pixel size in step and shoot mode. Each head acquired projection data at 20 angles during 8s. The frame duration including camera head movements was 4.1 minutes. The individual SPECT-frames were reconstructed into 128x128 matrices with Mediso's iterative Tera Tomo 3D ordered subset expectation maximization (OSEM) recon-struction including CT-based attenuation correction, Monte Carlo based (medium quality) scatter correction and resolution recovery. Regularization was set at "medium"; no pre- or post-filtering was applied and the number of updates (called iterations in the software) was 144 with 48 iterations and 3 subsets. The resulting isotropic voxel size was 1.8x1.8x1.8 mm. The SPECT-scanner with the described set-up was quantitatively calibrated for Tc-99m. Values in the reconstructed SPECT-images were therefore saved in Bq/ml.

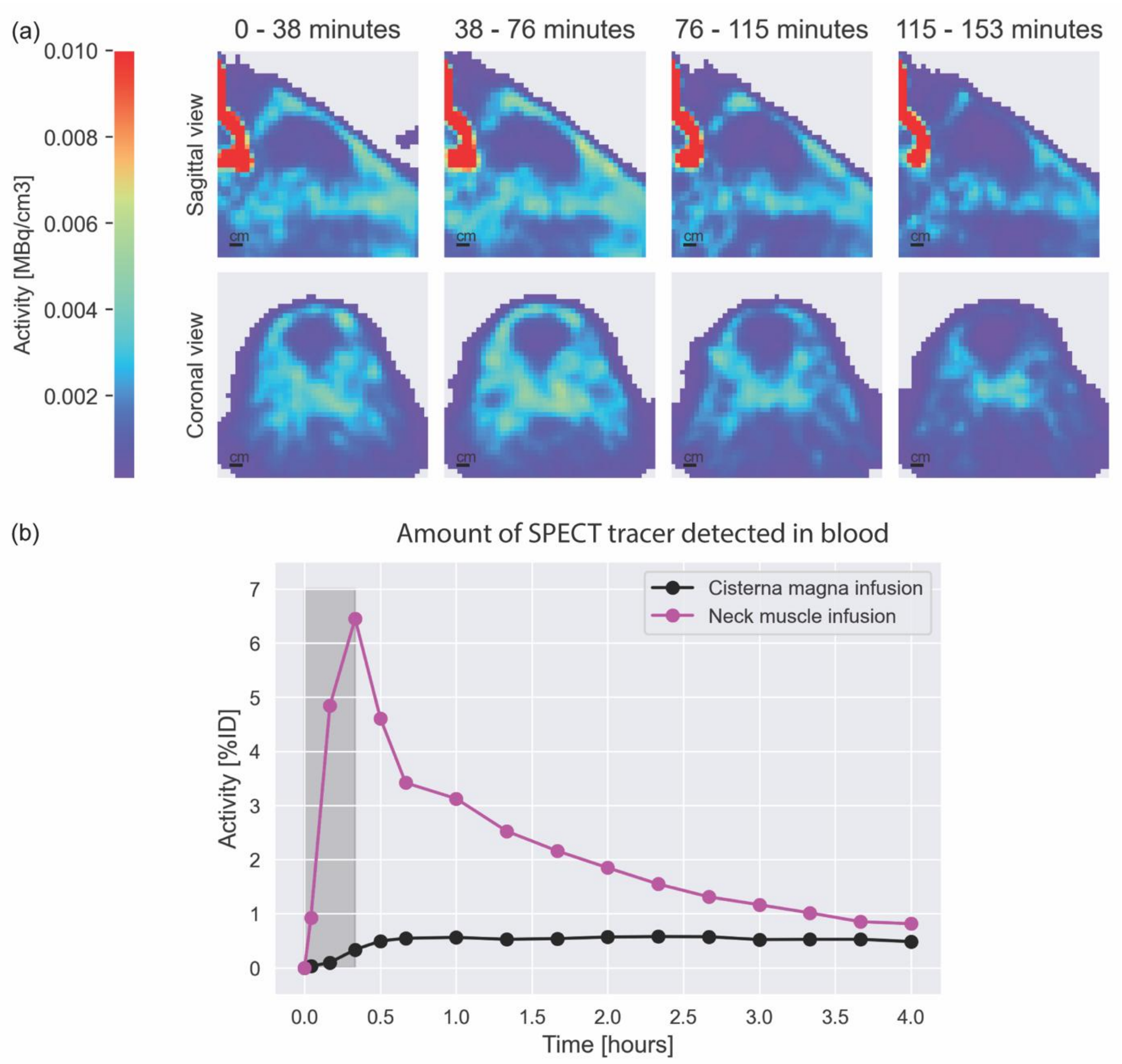


**Supplementary Figure 1.** *Unsuccessful CM Cannulation*. After initial successful cannula placement in the CM of the pig, the catheter was removed too far and the tip ended up remained in the neck muscles of the pig. A) Tracer accumulation in the neck muscles of the pig and no visible tracer distribution in the intracranial space. B) Tracer activity in the blood shows an initial fast increase and a subsequent drop over time. For comparison, tracer activity in the blood after the successful CM cannulation shows a general lower and timewise more constant distribution.

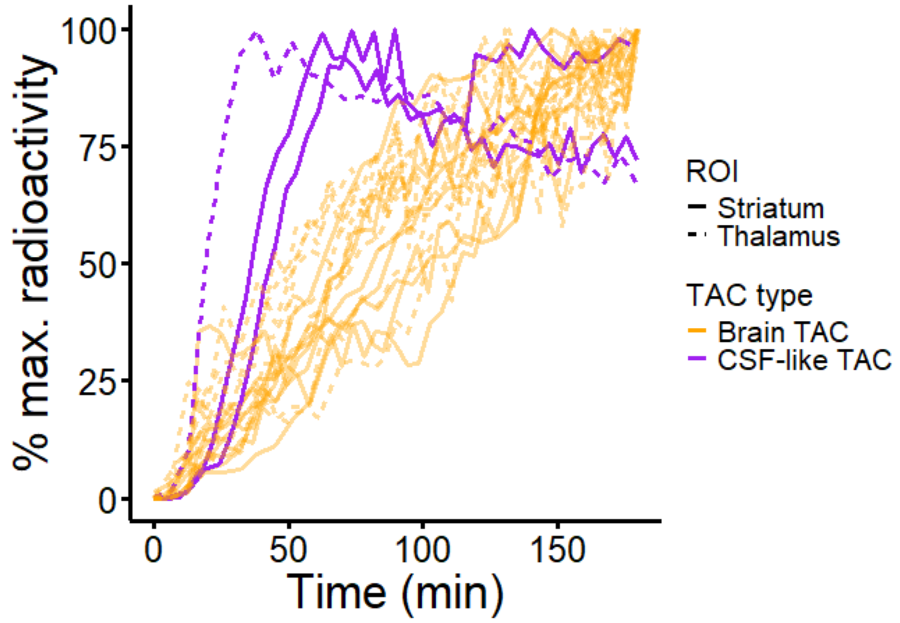


**Supplementary Figure 2. Striatum and thalamus TACs coloured by shape.** TACs are normalized to the maximum radioactivity for each pig to normalize across pigs. CSF-like TAC shape have a sharp increase in the beginning and decrease over time (in purple) while the brain-like TACs increase steadily over time (in orange). Striatum TACs are in continuous line while thalamus TAC are in discontinued line.
